# Expression of the long noncoding RNA DINO in HPV positive cervical cancer cells reactivates the dormant TP53 tumor suppressor through ATM/CHK2 signaling

**DOI:** 10.1101/2020.05.08.085555

**Authors:** Surendra Sharma, Karl Munger

## Abstract

Tumor cells overcome the cytostatic and cytotoxic restraints of TP53 tumor suppressor signaling through a variety of mechanisms. High-risk human papillomavirus (HPV) positive tumor cells retain wild type TP53 because the HPV E6/UBE3A ubiquitin ligase complex targets TP53 for proteasomal degradation. While restoration of TP53 in tumor cells holds great promise for cancer therapy, attempts to functionally restore the dormant TP53 tumor suppressor in HPV positive cancer cells by inhibiting the HPV E6/UBE3A ubiquitin ligase complex have not yet been successful. The Damage Induced long noncoding RNA, DINO, (DINOL) is a TP53 transcriptional target that has been reported to bind to and stabilize TP53, thereby amplifying TP53 signaling. We show that HPV positive cervical carcinoma cells contain low levels of DINO because of HPV E6/UBE3A mediated TP53 degradation. Acute DINO expression overrides HPV16 E6/UBE3A mediated TP53 degradation, causing TP53 stabilization and increased expression of TP53 transcriptional target genes. This causes a marked sensitization to chemotherapy agents and renders cells vulnerable to metabolic stress. Acute DINO expression in HPV positive cervical cancer cells induces hallmarks of DNA damage response signaling and TP53 activation involves ATM/CHK2 signaling. DINO upregulation in response to DNA damage is independent of ATM/CHK2 and can occur in cancer cells that express mutant TP53.

**IMPORTANCE:** Functional restoration of the TP53 tumor suppressor holds great promise for anti-cancer therapy. Current strategies are focused on modulating TP53 regulatory proteins. Long noncoding RNAs (lncRNAs) have emerged as important regulators of TP53 as well as modulators of downstream tumor suppressive transcriptional responses. Unlike many other cancer types, human papillomavirus (HPV) positive cancer cells retain wild type TP53 that is rendered dysfunctional by the viral E6 protein. We show that acute expression of the Damage Induced long Noncoding RNA, DINO, a known TP53 transcriptional target and functional modulator, causes TP53 reactivation in HPV positive cervical cancer cells. This causes increased vulnerability to standard chemotherapeutics as well as biguanide compounds that cause metabolic stress. Hence, strategies that target DINO may be useful for restoring TP53 tumor suppressor activity in HPV positive cancers and other tumor types that retain wild type TP53.

## INTRODUCTION

The productive viral life cycle of human papillomaviruses (HPVs) is confined to postmitotic, terminally differentiated squamous epithelial cells. Synthesis of viral progeny is critically dependent on cellular replication proteins. Hence, HPVs need to enforce continued expression of cellular proteins that are necessary for synthesis of viral genomes in terminally differentiated epithelial cells. High-risk HPV E7 proteins target the retinoblastoma tumor suppressor protein (RB1) for degradation (1, 2). This causes aberrant S-phase entry, which is sensed and leads to activation of the TP53 tumor suppressor protein. TP53 is a DNA binding transcription factor that is present at low levels in normal cells due to rapid proteasome mediated turnover by the MDM2 ubiquitin ligase (3–5). When activated, TP53 is stabilized, accumulates to high levels and engages cytostatic and cytotoxic transcriptional programs (6, 7). To blunt cell-abortive signals caused by HPV E7 expression, the high-risk HPV E6 proteins inactivate TP53 by forming a complex with the cellular ubiquitin ligase, UBE3A (E6-AP), thereby targeting TP53 for ubiquitination and proteasomal degradation (8–10). As a consequence of HPV E6/UBE3A mediated inactivation, most high-risk HPV associated tumors retain wild type TP53 (11, 12).

The TP53 tumor suppressor is mutated and rendered dysfunctional in most human cancers by mutation or by dysregulated expression of TP53 tumor suppressor pathway components (7). Restoration of TP53 signaling caused tumor regression in a variety of preclinical mouse models of human cancers, albeit through different mechanisms (13–18). This has prompted a keen interest in developing therapies aimed at restoring TP53 activity in cancers. Most of the pharmacological strategies that are currently tested in pre-clinical models or are in clinical trials target the MDM2-mediated TP53 degradation pathway (19–21). In HPV oncogene expressing cells there is a switch from MDM2-to HPV E6/UBE3A-mediated TP53 degradation (22). Reactivation of the dormant TP53 tumor suppressor in the HPV18 positive HeLa cervical cancer line by selectively repressing endogenous HPV18 E6 expression was shown to cause senescence and apoptosis (23). Hence, reactivation of TP53 tumor suppressor signaling may also have therapeutic benefits in HPV positive human tumors. Multiple strategies to hinder E6/UBE3A mediated TP53 inactivation have been explored, including inhibition of E6 or UBE3A expression (24, 25), and disruption of the TP53/E6/UBE3A complex with cellular proteins (26, 27), artificial E6 targeting microRNAs (28), RNA aptamers (29) synthetic peptides (30–32) as well as small molecules (33–35). However, none of these strategies has been translated to the clinic. Therefore, alternative approaches to reactivate the TP53 tumor suppressor pathway in HPV-associated cancers need to be explored.

The vast majority of the human transcriptome consists of noncoding RNAs. Several classes of non-coding RNAs, including microRNAs, circular RNAs and long noncoding RNAs (lncRNAs) regulate cellular processes that are dysregulated during carcinogenesis (36). Long noncoding RNAs (lncRNAs) denote a large family of RNAs of >200 nucleotides and with no or limited coding capacity of <100 amino acids. LncRNAs can form complexes with RNAs, DNAs and proteins and modulate cellular processes that have been referred to as hallmarks of human cancer (37, 38). Similarly, expression of the HPV E6 and E7 genes has been shown to remodel the cellular lncRNA transcriptome (39). Many lncRNAs have been identified as upstream and/or downstream components of the TP53 tumor suppressor pathway (40–42). The Damage Induced Noncoding (DINO) lncRNA is induced by TP53 in response to DNA damage. In addition, DINO has been reported to amplify the TP53 transcriptional response by binding and stabilizing TP53 and TP53/DINO complexes were detected bound to TP53 transcriptional response elements (43). Here we report that acute DINO expression at levels similar to those observed in response to DNA damage stabilizes TP53 and restores TP53 tumor suppressor activity in HPV positive cervical cancer cell lines. This causes increased sensitivity to standard-of-care chemotherapy agents and vulnerability to metabolic stress. Surprisingly, we found that, unlike the canonical TP53 target CDKN1A, DINO upregulation in response to DNA damage is through a pathway that is independent of ATM/CHK2 and was also observed in the HPV negative C33A cervical cancer cell line that expresses a DNA binding defective TP53 mutant. Moreover, acute DINO expression in HPV positive cervical cancer cells induces hallmarks of DNA damage signaling and activates TP53 through ATM/CHK2 signaling.

## RESULTS

### Low DINO levels in HPV-positive cervical cancer cells are a result of HPV E6-mediated TP53 degradation

We previously reported that DINO expression correlated with TP53 levels in HPV E6 and/or E7 expressing human foreskin keratinocytes (HFKs) (44). Given that HPV positive cervical cancer lines express E6 and E7 and retain wild-type TP53, we hypothesized that they may express DINO at similarly low levels as HPV16 E6/E7 expressing HFKs. To test this, we assessed DINO expression in HPV16 positive CaSki, HPV16 positive SiHa, and HPV18 positive HeLa cervical carcinoma lines by quantitative reverse transcription polymerase chain reaction (qRT-PCR) assays. As a control, we analyzed DINO expression in the HPV-negative C33A cervical carcinoma line that express a DNA binding-defective TP53 cancer hot-spot mutant where the arginine residue at position 273 is changed to a histidine (R273H) (45–47). These experiments showed that HPV positive cervical carcinoma lines expressed DINO at levels that were even lower than HPV16 E6/E7 expressing HFKs. The mutant TP53 expressing, HPV negative C33A cervical cancer cells contained even lower DINO levels (Figure 1A).

**Figure 1:**
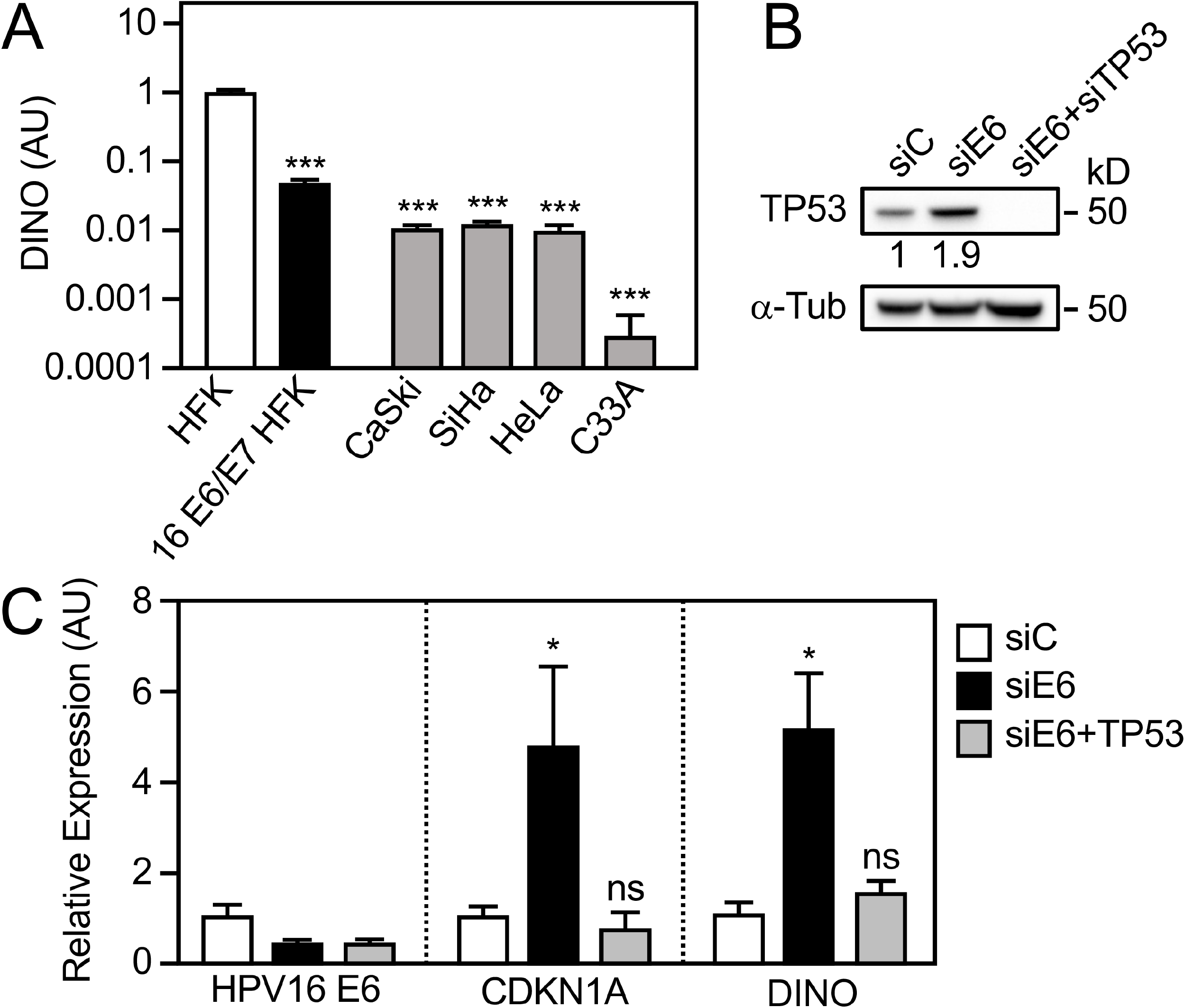
Low DINO levels in HPV-positive cervical cancer cells are a result of E6-mediated TP53 inactivation. DINO levels in HPV positive CaSki (HPV16), SiHa (HPV16), HeLa (HPV18) and HPV negative C33A cervical carcinoma cell lines were determined by quantitative reverse transcription polymerase chain reaction (qRT-PCR) assays. DINO levels in control vector transduced primary human foreskin keratinocytes (HFK) and HPV16 E6/E7 expressing HFKs (16 E6/E7 HFKs) are shown for comparison (A). SiHa cells were transfected with an siRNA targeting HPV16 E6 alone or in combination with TP53 targeting siRNAs. HPV16 E6 and TP53 depletion was validated by measuring TP53 protein levels by immunoblotting (B) and by quantifying E6 mRNA levels qRT-PCR (C). The TP53 dependent effects of HPV16 E6 depletion on DINO and the canonical TP53 transcriptional target, CDKN1A were also assessed by qRT-PCR (C). Expression data is presented in arbitrary units (AU) and is normalized to expression of the RPLP0 housekeeping gene. The positions of marker proteins with their apparent molecular weights in kilodaltons (kD) are indicated with the Western blots. Bar graphs represent means ± SEM (n=3). *** p < 0.001, * p < 0.05 and ns = non-significant (Student’s t-test).

To determine whether the low DINO levels in HPV positive cervical cancer lines were a consequence of HPV E6/UBE3A-mediated TP53 destabilization, HPV16 E6, alone or in combination with TP53, was depleted in HPV16 positive SiHa cells by transient transfection of the corresponding siRNAs. To assess the efficiency of HPV16 E6 and TP53 depletion, TP53 protein levels were assessed by western blotting. As expected, HPV16 E6 depletion caused an increase in TP53 steady state levels, which was abrogated by TP53 co-depletion (Figure 1B). Like the canonical TP53 transcriptional target, CDKN1A, DINO levels increased upon E6 depletion and this effect was abrogated by co-depletion of TP53 (Figure 1C). Hence, the low levels of DINO in HPV positive cervical carcinoma lines likely represent a consequence of E6/UBE3A mediated TP53 destabilization.

### Acute DINO expression in HPV-positive cervical cancer cells reconstitutes dormant TP53 tumor suppressor activity

DINO expression is regulated by TP53 and has been reported to bind and stabilize TP53, thereby amplifying TP53 signaling. We have previously shown that HPV16 E7 expression causes TP53 stabilization and activation through DINO (44). Given that HPV16 E6 depletion increased DINO levels and caused a TP53 dependent increase in the TP53 transcriptional target CDKN1A in the HPV positive SiHa cervical cancer line (Figure 1), we next determined whether the dormant TP53 tumor suppressor pathway may be restored by DINO expression. Because high-level ectopic DINO expression may trigger TP53 dependent cytotoxic and/or cytostatic responses, we created vectors for doxycycline regulated DINO expression and generated HPV16 positive SiHa and CaSki cervical cancer cell populations with doxycycline regulated DINO expression. Cells expressing a vector with doxycycline inducible green fluorescent protein (GFP) expression were also made to be used as controls. To ensure that doxycycline induced DINO expression by this method mimic DINO induction by a biologically relevant stimulus, we compared SiHa cells with doxycycline induced DINO expression to DINO expression in response to DNA damage. The chemotherapy agent, doxorubicin, a known, potent inducer of DINO expression (43) was used for these experiments. Doxycycline induction caused a similar increase in DINO expression as treatment with doxorubicin (Figure 2A). Moreover, subcellular fractionation experiments revealed that increases in cytoplasmic and nuclear DINO (Figures 2B, C) were similar in doxycycline induced and doxorubicin treated SiHa cells. Hence, doxycycline mediated DINO expression closely mirrors DINO induction in response to DNA damage.

**Figure 2:**
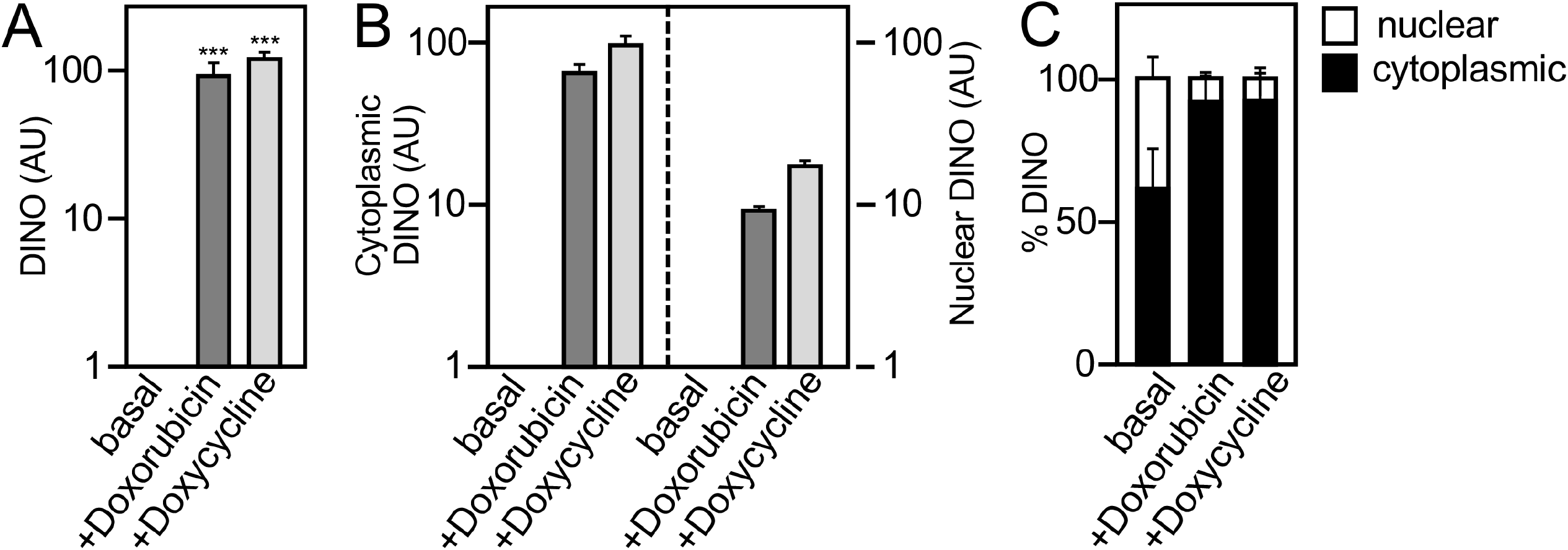
Doxycycline mediated DINO expression mimics induction by DNA damage. DINO expression as analyzed by qRT-PCR in control vector transduced SiHa cells (basal) or treated with 0.2 μg/ml doxorubicin for 24 hours (+Doxorubicin) as compared to acute DINO expression by treating inducible DINO vector transduced SiHa cells with 1 μg/ml doxycycline for 48 hours (+Doxycycline) (A). Quantification of the increases in the cytoplasmic and nuclear DINO levels by qRT-PCR (B). Assessment of the relative increases in the nuclear and cytoplasmic DINO pools by qRT-PCR (C). Expression data are presented in arbitrary units (AU) and are normalized to expression of the RPLP0 housekeeping gene. Bar graphs represent means ± SEM (n=3). *** p < 0.001 (Student’s t-test).

After validating the doxycycline mediated expression system, we tested whether doxycycline induced, acute DINO expression may override HPV16 E6/UBE3A mediated TP53 inactivation and restore TP53 levels and/or activity in the HPV16 positive SiHa (Figure 3A) and CaSki (Figure 3B) cervical cancer cell lines. DINO expression was validated by qRT-PCR assays (Figures 3A and 3B, left panels). Immunoblot experiments revealed higher levels of TP53 and concomitant increased expression of the canonical TP53 transcriptional target, CDKN1A, in SiHa and CaSki cells in response to DINO expression (Figure 3A and 3B, right panels). These results show that acute DINO expression causes functional reactivation of dormant TP53 tumor suppressor signaling in HPV-positive cervical carcinoma lines.

**Figure 3:**
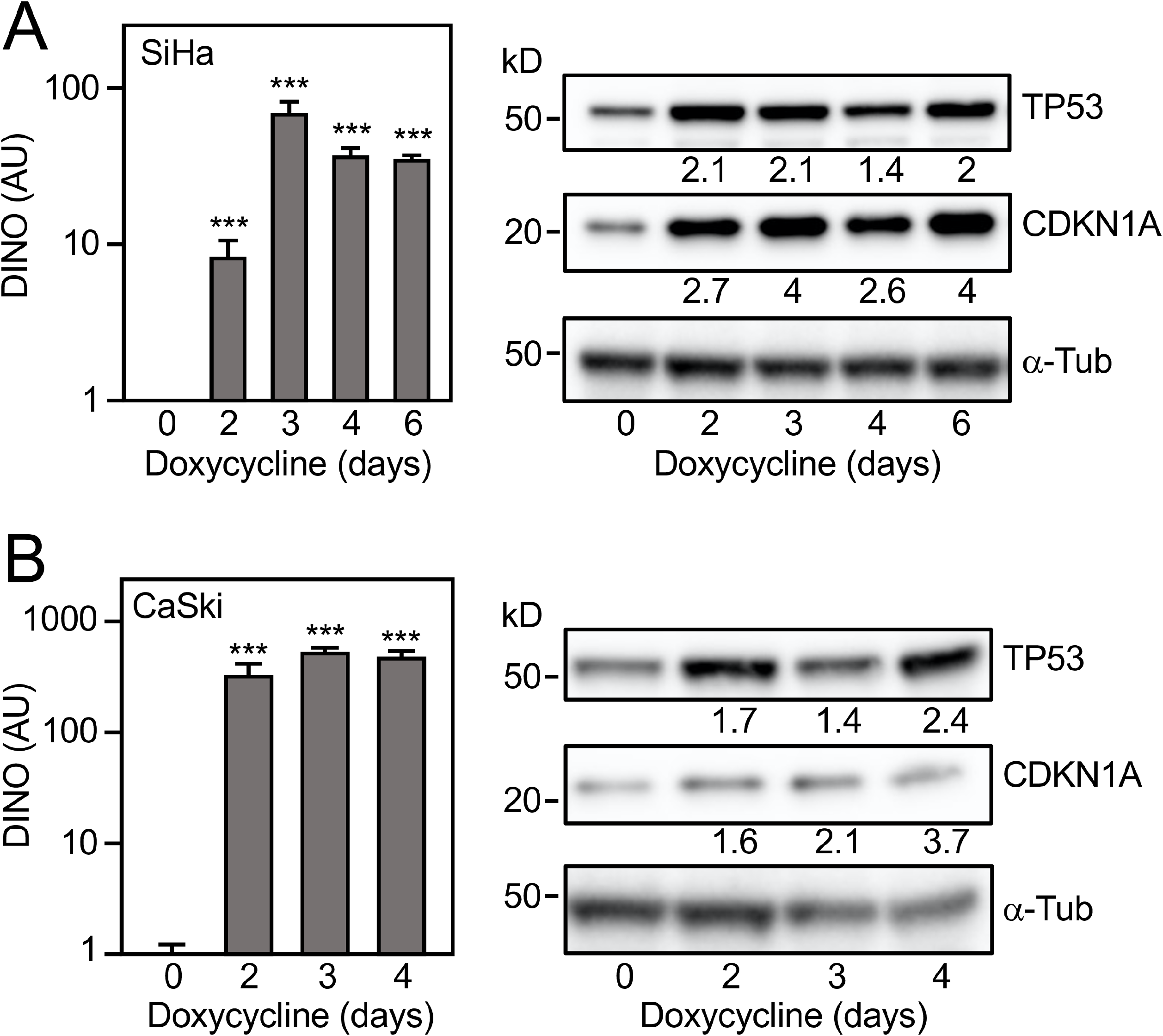
Acute DINO expression in HPV positive cervical cancer cells causes reactivation TP53 signaling. DINO expression in inducible DINO vector transduced HPV16 positive SiHa (A) and CaSki (B) cervical cancer cells after treating with 1 μg/ml doxycycline for the indicated number of days as determined by qRT-PCR. Expression data are presented in arbitrary units (AU) and are normalized to expression of the RPLP0 housekeeping gene. Bar graphs represent means ± SEM (n=3). *** p < 0.001 and * p < 0.05 (Student’s t-test) (left panels). Changes in TP53 and CDKN1A proteins levels were measured by immunoblotting with α-tubulin (α-Tub) serving as a loading control (right panels). The positions of marker proteins with their apparent molecular weights in kilodaltons (kD) are indicated.

### DINO overrides HPV16 E6/UBE3A mediated TP53 degradation

DINO was shown to stabilize TP53 in normal human cells where TP53 turnover is governed by MDM2 mediated ubiquitination (43). To determine whether the observed increases in TP53 upon DINO expression are caused by overriding HPV E6/UBE3A mediated TP53 turnover, we performed a cycloheximide chase experiment. After inducing DINO expression in SiHa cells by a 48-hour doxycycline treatment, cycloheximide was added to inhibit new protein synthesis. SiHa cell with conditional GFP expression were used as controls. TP53 steady state levels were determined in extracts harvested at different times after cycloheximide treatment (Figure 4, top panel). Quantification revealed that TP53 half-life was increased from ∼30 minutes in GFP expressing control cells to ∼45 minutes in DINO expressing SiHa cells (Figure 4, bottom panel). Hence acute DINO expression can override HPV16 E6/UBE3A mediated TP53 degradation.

**Figure 4:**
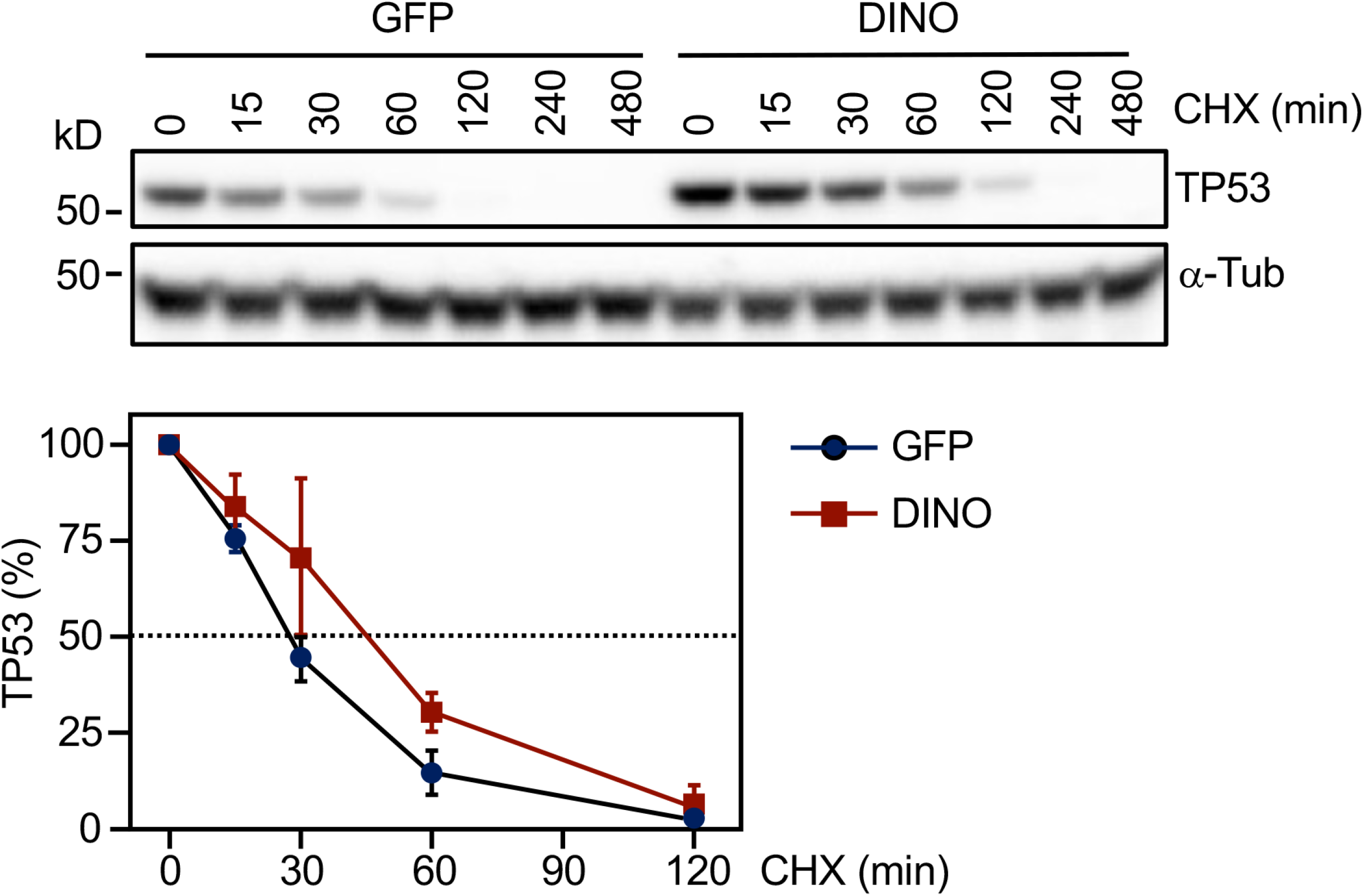
DINO expression increases TP53 protein stability. HPV16 positive SiHa cervical cancer cells transduced with inducible DINO or GFP control vector were treated with 1 μg/ml doxycycline for 48 hours, followed by 100 μg/ml cycloheximide (CHX) to block protein synthesis. Proteins lysates were prepared at various time points after cycloheximide addition and TP53 levels were measured by immunoblotting and quantified to estimate the half-life of TP53 with α-tubulin (α-Tub) serving as a loading control. A representative blot is shown in the top panel. The positions of marker proteins with their apparent molecular weights in kilodaltons (kD) are indicated. Quantification representing using means ± SEM of three experiments is shown in the bottom panel.

### Ectopic DINO increases the susceptibility of HPV positive cervical cancer cell lines to inducer of metabolic stress

We previously reported that HPV16 E7 expressing HFKs have elevated DINO and that DINO depletion protected these cells from metabolic stress induced by starvation or by treatment with the antidiabetic biguanide drug metformin (44). Metformin and related biguanides are antidiabetic drugs that are investigated for potential repurposing as anticancer agents because they activate AMP-activated protein kinase, which in turn has been reported to activate TP53 (48–50). Hence, we tested whether acute DINO expression in HPV positive cancer cells with low endogenous DINO expression may render them vulnerable to metabolic stress. As expected, doxycycline induction of DINO in CaSki cells triggered increased mRNA expression of the TP53 transcriptional target CDKN1A and elevated TP53 steady state levels (Figure 5A). CaSki cells with acute DINO overexpression were significantly more vulnerable to serum deprivation and exhibited significantly enhanced sensitivity to metformin and the related more potent biguanide drug, phenformin (Figure 5B). These experiments revealed that, consistent with our previous study (44), DINO is a crucial modulator of cell death during metabolic stress.

**Figure 5:**
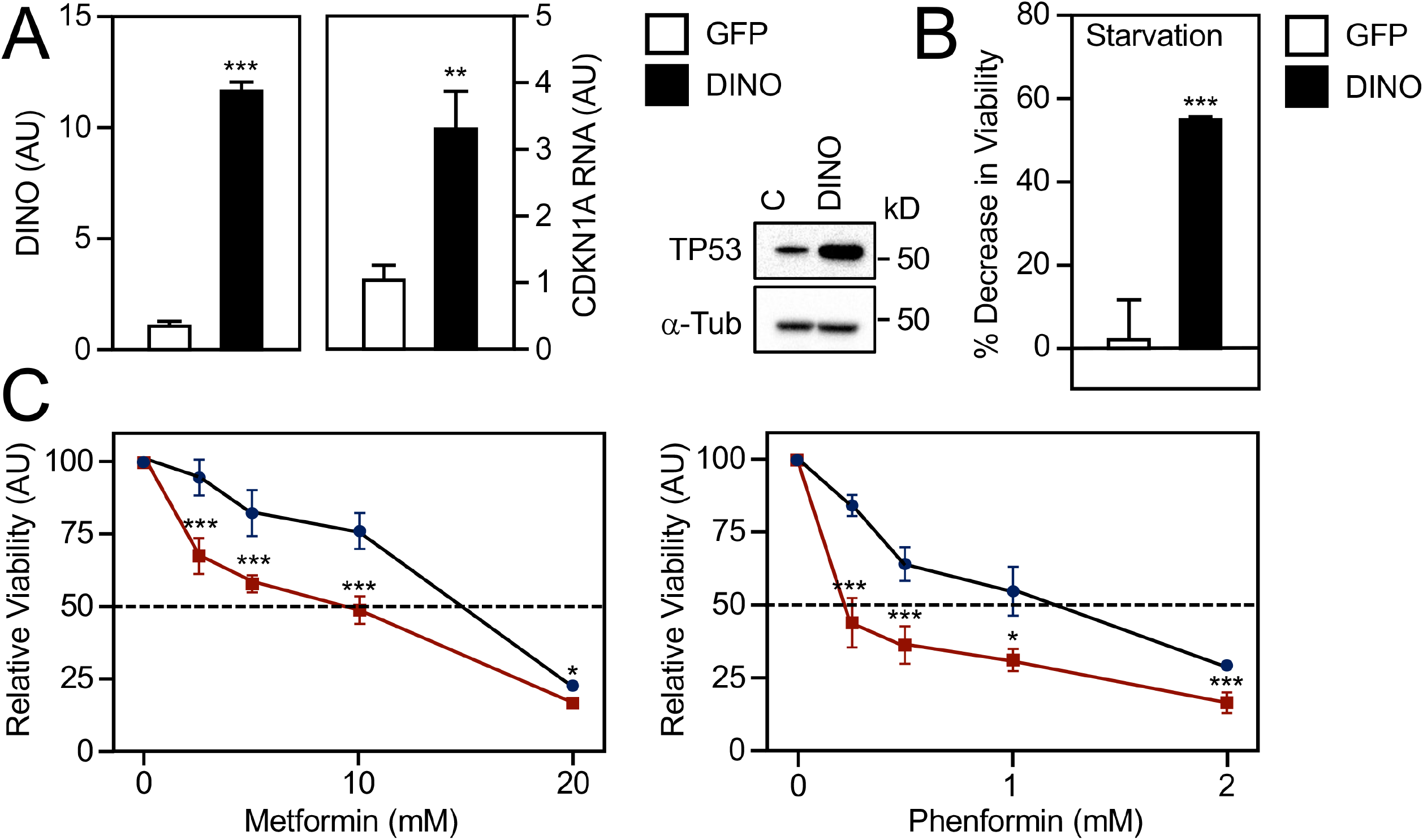
Acute DINO expression increases susceptibility to metabolic stress. HPV16 positive CaSki cervical cancer cells transduced with inducible DINO or GFP control vector were each treated with 1 μg/ml doxycycline for 48 hours. DINO expression and mRNA levels of the TP53 transcriptional target, CDKN1A, were determined by qRT-PCR. Expression data are presented in arbitrary units (AU) and are normalized to expression of the RPLP0 housekeeping gene (A-left panel). Concomitant changes in TP53 proteins levels were assessed by immunoblotting blotting with α-tubulin (α-Tub) serving as a loading control. The positions of marker proteins with their apparent molecular weights in kilodaltons (kD) are indicated (A-right panel). GFP or DINO expressing CaSki cells were incubated with serum free medium (B) or grown in normal serum containing medium supplemented with the indicated concentrations of metformin or phenformin for 72 hours (C), and cell viability was assessed. Graphs represent means ± SEM (n=3). *** p < 0.001, ** p < 0.01, * p < 0.05 (Student’s t-test).

### Ectopic DINO increases susceptibility of HPV positive cervical cancer cell lines to chemotherapeutics

The lack of functional TP53 likely renders HPV-positive cervical tumor refractory to standard chemotherapy and radiation treatments (51). Hence, we tested whether acute, ectopic DINO expression may sensitize HPV positive cervical cancer cells to chemotherapy drugs. We used CaSki cells for these experiments because SiHa cells were shown to be highly resistant to chemotherapy drugs (52)Chemotherapy involving the alkylating agent cisplatin and related platinum compounds have long been the standard of care for advanced, recurrent cervical cancers (53). Other widely used anti-cancer chemotherapy agents include the antimetabolite 5-fluorouracil (5-FU), the alkylating agent mitomycin C, and the DNA intercalating agent doxorubicin. Acute, doxycycline mediated DINO expression rendered CaSki markedly more sensitive to cisplatin (Figure 6A), doxorubicin (Figure 6B), mitomycin C (Figure 6C), and 5-FU (Figure 6D) as compared to control cells with doxycycline regulated GFP expression. Hence the sensitivity of cervical cancer cells to a variety of chemotherapy agents is modulated by DINO and presumably represents a consequence of reactivating the dormant TP53 tumor suppressor pathway.

**Figure 6:**
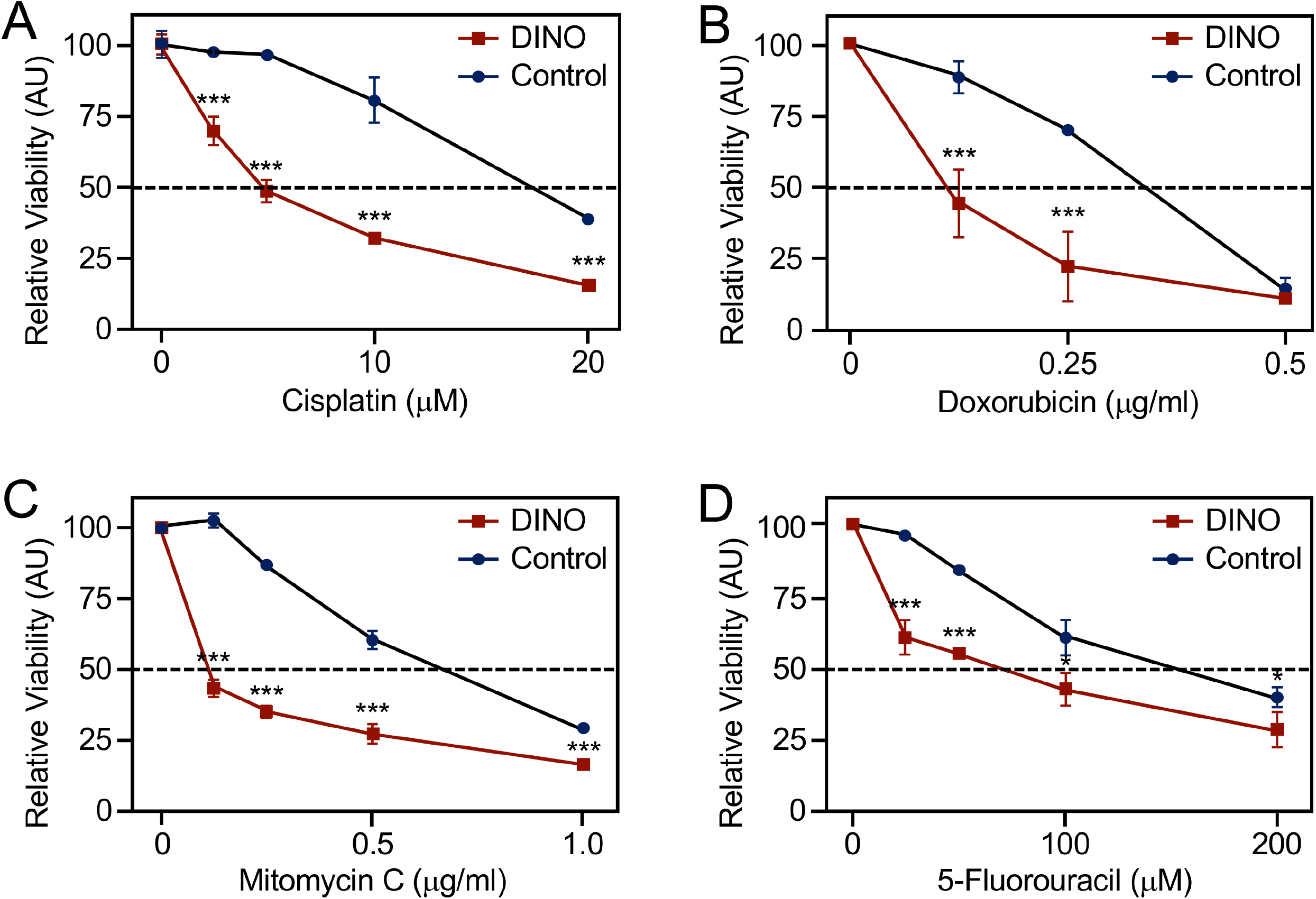
Acute DINO expression increases susceptibility to chemotherapy agents. Inducible DINO vector transduced, HPV16 positive CaSki cervical cancer cells were treated with the indicated concentrations of cisplatin (A), doxorubicin (B), mitomycin C (C) and 5-fluorouracil (D). Cell viability was assessed at 72 hours post treatment. Graphs represent means ± SEM from single representative experiments, each performed in triplicate. *** p < 0.001, * p < 0.05 (Student’s t-test). Similar results were obtained in three independent experiments.

### DINO induction by DNA damage is independent of the ATM/CHK2/TP53 DNA damage signaling pathway

Increased DINO expression in response to DNA damage has been reported to be TP53 dependent (43). To discern whether DNA damage mediated DINO expression is also TP53 dependent in cervical carcinoma lines, we performed epistasis analysis using small molecule inhibitors of ataxia-telangiectasia mutated (ATM) and checkpoint kinase 2 (CHK2), two kinases that are well known to signal DNA damage to TP53 (54, 55). As expected, doxorubicin treatment of SiHa and CaSki cells caused a robust increase of DINO as well as the TP53 transcriptional target, CDKN1A (Figure 7A, right and left panels, respectively). Consistent with efficient inhibition of DNA damage mediated TP53 activation by ATM or CHK2 inhibition, expression of the canonical TP53 target gene, CDKN1A, was dramatically reduced. In contrast however, ATM or CHK2 inhibition did not significantly interfere with induction of DINO expression by doxorubicin in either SiHa or CaSki cells (Figure 7A). These results indicate that DINO induction by doxorubicin is less dependent on ATM/CHK2 mediated TP53 activation than CDKN1A.

**Figure 7:**
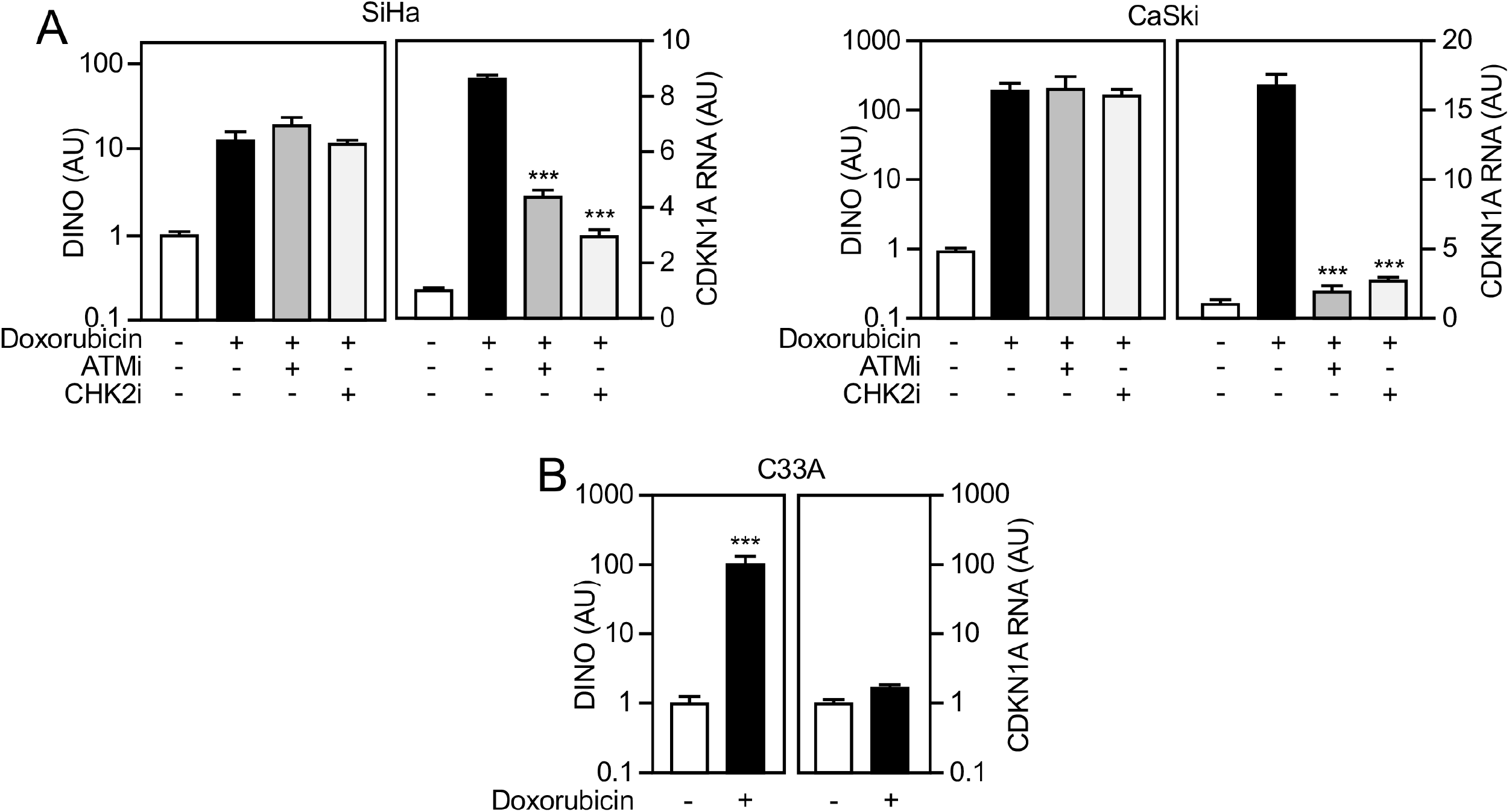
DINO induction by DNA damage can occur independent of ATM/CHK2/TP53 DNA damage signaling. The HPV16 positive SiHa (left) and CaSki (right) cervical cancer lines were treated with 0.2 μg/ml doxorubicin and either 10 μM of the ATM inhibitor KU-55933 (ATMi), or 5 μM of the CHK2 inhibitor BML-277 (CHK2i). After 24 hours, expression of DINO and the canonical TP53 target gene CDKN1A was determined by qRT-PCR (A). Doxorubicin-mediated induction of DINO and CDKN1A mRNA was also assessed in the HPV negative C33A cell line that expresses the DNA binding defective, TP53 R273H cancer hotspot mutant (B). Expression data are presented in arbitrary units (AU) and are normalized to expression of the RPLP0 housekeeping gene. Bar graphs represent means ± SEM (n=3). *** p < 0.001 (Student’s t-test).

Based on this finding we next tested whether doxorubicin treatment may also trigger DINO expression in the HPV-negative C33A cervical carcinoma line which expresses the DNA binding defective TP53 R273H mutant (45–47). Doxorubicin treatment caused a robust increase in DINO levels whereas expression of the TP53 target gene, CDKN1A, remained unaltered (Figure 7B). Hence, DINO induction in response doxorubicin can efficiently occur in cells that lack functional TP53.

### Acute DINO expression induces hallmarks of DNA damage signaling

DINO has been reported to bind TP53 thereby stabilizing TP53 and amplifying DNA damage signaling (43). This predicts that acute DINO expression will cause TP53 stabilization independent of upstream DNA damage signaling. To experimentally test this model, we assessed ATM and CHK2 phosphorylation as well as TP53 stabilization and induction of TP53 transcriptional targets in response to acute, doxycycline mediated DINO expression in CaSki and SiHa cervical carcinoma cells (Figure 8A). Surprisingly, we detected robust phosphorylation of CHK2 at threonine 68 and ATM phosphorylation at serine 1981 that paralleled TP53 stabilization and CDKN1A protein expression in both cell lines. In SiHa but not in CaSki cells, we also observed induction of the p90 and p75 MDM2 isoforms with a concomitant decrease in TP53 levels and CDKN1A expression (Figure 8A). These results show that acute DINO expression triggers ATM and CHK2 activation. To determine whether acute DINO expression also induces other hallmarks of the DNA damage response, we investigated the appearance of 53BP1 nuclear foci in CaSki cells. Doxycycline treated cells with conditional GFP expression were used as negative controls. Acute expression of DINO caused an increased appearance of 53BP1 nuclear foci (Figure 8B).

**Figure 8:**
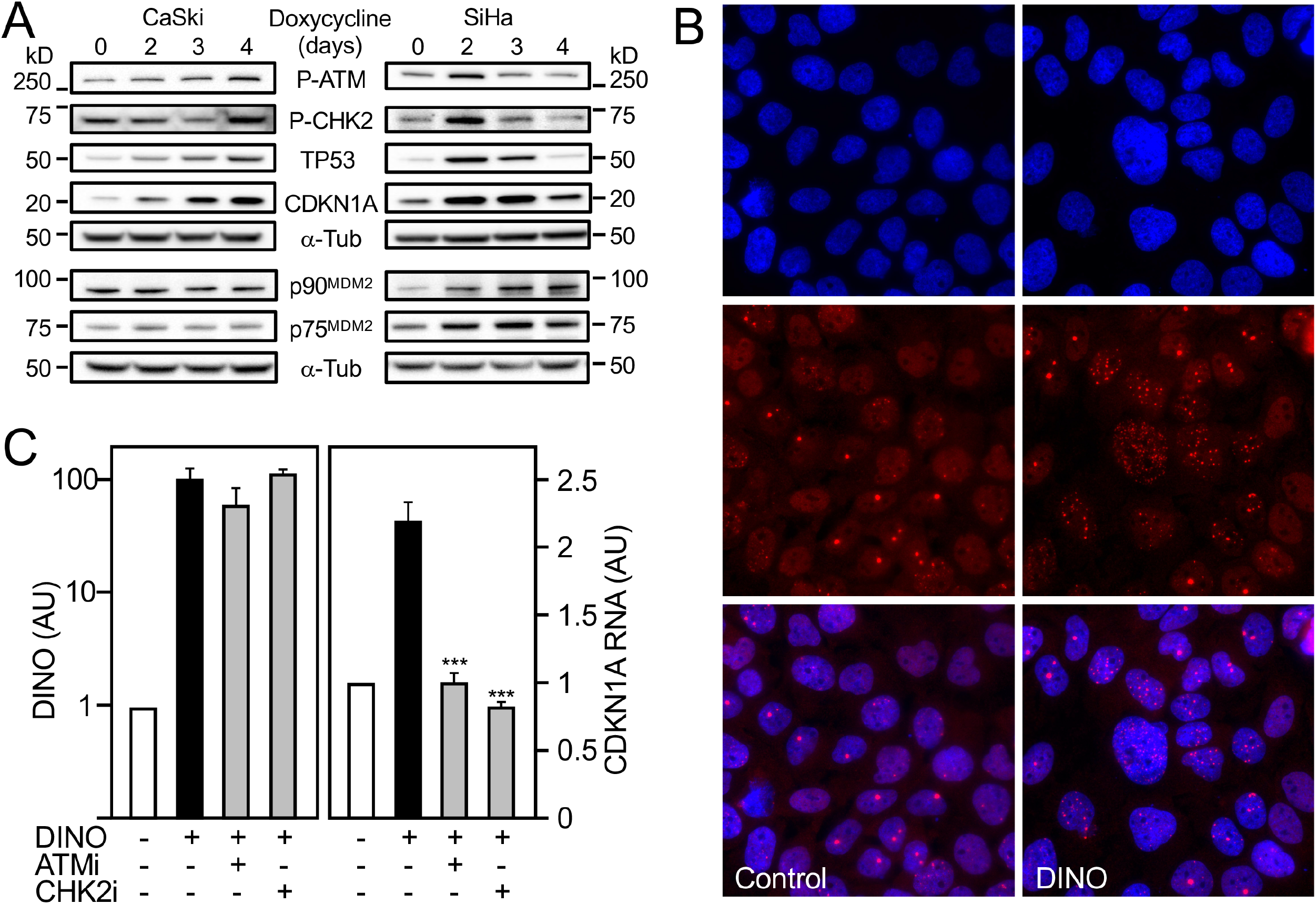
Acute DINO expression induces hallmarks of DNA damage signaling. HPV16 positive CaSki (left) and SiHa (right) cervical cancer lines with inducible DINO expression were treated with 1 μg/ml doxycycline and levels of phospho-Ser1981 ATM (P-ATM), phospho-Thr68 CHK2 (P-CHK2), TP53, CDKN1A, and the p90 and p75 MDM2 isoforms were assessed by immunoblot analysis with α-tubulin (α-Tub) serving as a loading control. The MDM2 blots were performed in a separate experiment but using aliquots of the same protein extracts as those used for the other blots. The positions of marker proteins with their apparent molecular weights in kilodaltons (kD) are indicated (A). Nuclear 53BP1 foci in CaSki cells in control and after doxycycline mediated DINO expression. Nuclei were stained with DAPI. Representative immunofluorescence microscopy images are shown; similar results were obtained in two independently performed microscopy experiments (B). CaSki cells with inducible DINO expression were treated with 1 μg/ml doxycycline and 10 μM of the ATM inhibitor KU-55933 (ATMi) or 5 μM of the CHK2 inhibitor BML-277 (CHK2i) and expression of DINO and the canonical transcriptional TP53 target gene CDKN1A were determined by qRT-PCR. Expression data are presented in arbitrary units (AU) and are normalized to expression of the RPLP0 housekeeping gene. Bar graphs represent means ± SEM (n=3). *** p < 0.001 (Student’s t-test) (C).

To determine whether TP53 activation in response to acute DINO expression was dependent on ATM and CHK2 activation, we treated CaSki cells with CHK2 or ATM inhibitors and assessed mRNA expression of the canonical TP53 transcriptional target CDKN1A. While ATM and CHK2 inhibition did not interfere with doxycycline mediated DINO induction, these treatments blocked TP53 mediated CDKN1A mRNA expression (Figure 8C).

Taken together, our results indicate that DINO mediated TP53 stabilization and activation in cervical carcinoma cells is through a pathway that involves activation of the DNA damage signaling pathway.

## DISCUSSION

The TP53 tumor suppressor, dubbed the “guardian of the genome,” is a key component of an innate cellular defense signaling network that functions to remove cells that have suffered genetic or epigenetic alterations that interfere with normal cellular homeostasis, from the proliferative pool (6). Hence, acquisition of defects in the TP53 surveillance pathway is an almost universal and rate limiting step in human carcinogenesis. Functional restoration of TP53 tumor suppressor signaling in human cancers was shown to have cytostatic or cytotoxic effects (56). In addition to proteins, non-coding RNAs particularly microRNAs and lncRNAs have emerged as key regulators and modulators of TP53 signaling (40–42, 57).

DINO is a lncRNA that functions as a regulator as well as a modulator of TP53 (43). DINO was identified as a TP53 transcriptional target that can amplify TP53 signaling by binding and stabilizing TP53 and was shown to be a component of DNA bound TP53 tetramers. DINO binding involves the C-terminal 31 amino acids of TP53 whereas MDM2, the ubiquitin ligase that targets TP53 for rapid proteasomal degradation, binds to TP53 sequences within the amino terminal transactivation domain (43, 58). Hence, it is unlikely that DINO and MDM2 directly compete for binding to the same TP53 sequences. Similarly, the TP53 C-terminal 31 amino acids are not necessary for E6/UBE3A binding, and, hence, interaction with DINO is not predicted to directly compete with E6/UBE3A binding (59). Interestingly, TP53 has long been known to interact with RNAs, and the TP53 C-terminus has been reported to be critical for RNA binding, which is regulated by the acetylation status of multiple lysine residues (60). It will be interesting to determine whether DINO binding is similarly regulated by TP53 post-translational modifications.

In this study, we show that HPV positive cervical cancer lines contain low levels of DINO as a consequence of HPV16 E6/UBE3A mediated degradation. Acute DINO expression that mimics DINO induction in response to DNA damage, causes TP53 stabilization and functional reactivation as evidenced by increased expression of the canonical TP53 transcriptional target, CDKN1A. In the case of SiHa cells, we consistently observed a dip in the increase of TP53 and CDKN1A protein levels at days 3 or 4 post doxycycline induced DINO expression. TP53 levels and activity are regulated by a positive and negative feedback loops (61–63). The observed transient TP53 decrease may indeed be caused by increased expression of the MDM2 p90 and p75 isoforms (64), which we observed in SiHa but less so in CaSki cells within the analyzed time course. Alternatively, given that these experiments were performed with polyclonal cell populations, the observed dip might be attributable to the heterogeneity of DINO induction levels. Nonetheless, the observed ∼2-fold increase in TP53 levels in response to acute DINO expression is similar to what we observed upon HPV16 E6 silencing.

Despite multiple attempts, we did not succeed in generating cervical cancer lines with constitutive, ectopic, high-level DINO expression. Hence, we generated cervical cancer lines with doxycycline regulated DINO expression. Doxycycline treatment increased DINO to similar levels as in cells that were treated with the DNA damage inducing chemotherapy agent, doxorubicin. Under both conditions, there is more dramatic increase of DINO in the cytoplasm than in the nucleus. The biological significance of these two DINO populations is currently unknown. It is noted, however, that TP53 shuttles between the nucleus and cytoplasm and that nuclear export is necessary for degradation by MDM2 and E6/UBE3A (65). Therefore, nuclear and cytoplasmic DINO could each modulate TP53 turnover and activity.

We were initially surprised that despite TP53 stabilization and functional reactivation, acute, doxycycline mediated induction of DINO did not noticeably inhibit the viability of SiHa and CaSki cells. This suggests that DINO expression achieved by doxycycline induction may not reach levels that surpass the critical threshold required to drive a TP53 mediated cytotoxic response (66). Importantly, however, TP53 mediated transcriptional responses also include expression of negative regulatory proteins, including the MDM2 ubiquitin ligase, that dampen TP53 activity. We consistently observed increased MDM2 protein expression in SiHa upon acute DINO expression cells with a concomitant decrease in TP53 and CDKN1A levels. Moreover, HPV16 E7 expressing keratinocytes express high levels of DINO and TP53 and they proliferate at a similar rate than parental or HPV16 E6/E7 co-expressing cells, which express low TP53 and DINO (44). HPV16 E7 expressing cells, however, are susceptible to cell death when subjected to metabolic stress (2). The vulnerability to metabolic stress was shown to be TP53 dependent (67) and abrogated upon DINO depletion (44). Here we show that that acute DINO expression markedly increases the vulnerability of cervical carcinoma cells to a variety of inducers of metabolic stress and increases their susceptibility to multiple classes of DNA damage inducing chemotherapy agents, including cis-platin which has long been the standard-of-care chemotherapy agent for the treatment of invasive cervical carcinoma.

After DNA damage, DINO was found in complex with TP53 and detected in DNA bound TP53 transcription factor complexes thereby amplifying TP53 tumor suppressor signaling (43). We were thus surprised that even after DNA damage, the major fraction of DINO was detected in the cytoplasm. Hence, DINO may have other cellular targets and possess biological activities that are independent of transcriptional, nuclear TP53 tumor suppressor signaling.

In support of this model, we discovered that in contrast to other TP53 responsive genes, induction of DINO by doxorubicin-mediated DNA damage occurs even when the CHK2 and ATM DNA damage kinases are inhibited. Moreover, DINO is induced by DNA damage in C33A cells that express the DNA binding defective TP53 R273H cancer hotspot mutant suggesting that DINO can be induced through additional ATM/CHK2 and TP53 independent mechanisms, although the molecular consequences of increased DINO expression in TP53 mutant cells are currently unknown. In addition to ATM/CHK2, DNA damage may also lead to activation of the ATM-and Rad3-related (ATR) kinase and/or the DNA-dependent protein kinase (DNA-PK) pathways, which, together with ATM, comprise “the trinity of kinases” that signal the DNA damage response (68). TP53 independent, epigenetic, mechanisms of DINO expression have also been described. The initial trigger of HPV16 E7 mediated DINO induction that leads to TP53 stabilization and activation involves the H3K27 demethylase KDM6A (44) and HPV16 E7 is well known to cause double strand DNA breaks (69, 70). Similarly, an orphan nuclear receptor, Nuclear Receptor Subfamily 2 Group E Member 3 (NR2E3) and the associated KDM1A (LSD1) H3K4 demethylase were shown to be necessary for efficient induction of DINO and TP53 activity in response to liver damage by N-acetyl-p-benzoquinone imine mediated oxidative stress (71). It is noted that in addition to DNA damage, doxorubicin can also induce free radical release thereby causing oxidative stress (72).

Our studies also provide evidence that in addition to TP53 binding, stabilization and by functioning as a component of DNA bound TP53 transcription factor complexes (43), DINO can cause or amplify TP53 activation through a mechanism that involves activation of DNA damage response signaling as evidenced by ATM and ATR activation and the appearance of cells that contain larger numbers of 53BP1 nuclear foci. Moreover, ATM or CHK2 inhibition interfered with DINO mediated TP53 activation as assessed by CDKN1A transcription.

LncRNAs have emerged as critical regulators of cellular DNA damage responses but the molecular mechanisms by which DINO drives DNA damage signaling remain to be determined. Similar to DINO, the long intergenic radiation-responsive noncoding RNAs (LIRRs) induce DNA damage signaling through mechanisms that are not clear (73). On the other hand, the GUARDIN lncRNA, which is essential for genomic stability, inhibits DNA damage through mechanisms involving the stabilization of telomeric repeat-binding factor 2 (TRF2) and Breast cancer type 1 susceptibility protein (BRCA1) (74). One possibility is that DINO expression induces double-stranded DNA breaks. Alternatively, DINO may drive the DNA damage response by modulating signaling events by scaffolding and recruiting critical proteins such as ATM, 53BP1, H2A.X and RNF168. Lastly, it is possible that the effect is indirect, i.e. that DINO regulates expression of other, coding or non-coding, genes that may induce DNA damage, modulate DNA damage repair, or DNA damage repair signaling. The HITT (HIF-1α inhibitor at translation level) lncRNA, for example, was reported to blunt ATM activation by directly binding to ATM and blocking recruitment of the MRE11-RAD50-NBS1 complex (75).

Taken together, these results indicate that DINO is not only a “Damage Induced Noncoding” RNA but may also function as a “Damage Inducing Noncoding RNA”. This hypothesis is based on results obtained with cells that were engineered to acutely express DINO in response to doxycycline. While this represent an artificial method of modulating DINO expression, it is noted that the overall increase in DINO expression and the subcellular DINO localization mirrors what is observed upon induction of DNA damage and hence, it is unlikely to be entirely superphysiological. Moreover, it is widely recognized that lncRNAs can act as components of DNA damage response and repair processes (76). It will be important to determine whether DINO directly induces DNA damage, whether it is a component of complexes that form to sense or mark sites of DNA damage, and/or are involved in DNA break repair.

In summary, we show that the dormant TP53 tumor suppressor pathway in HPV positive cervical cancer lines can be functionally reactivated by modulating DINO. Our results support a model whereby DINO activates TP53 through a CHK2/ATM dependent pathway. The concomitant ATM/CHK mediated activation of TP53 then triggers a further, TP53 dependent increase in DINO, which in turn amplifies DNA damage repair signaling. This causes further TP53 activation. DINO may also augment TP53 signaling as a component of DNA bound TP53 transcription factor complexes (43) (Figure 9). DINO mediated TP53 reactivation causes cervical cancer cells to be more sensitive to standard chemotherapeutics and induces marked vulnerability to metabolic stress. Moreover, we found that DNA damage can induce DINO expression despite ATM/CHK2 inhibition and in C33A cells that express the DNA binding defective TP53 R273H cancer hot spot mutant. These findings warrant further studies examining whether these DINO activities are also operative in non-HPV-associated tumors with different aberrations in the TP53 tumor suppressor pathway. Given that lncRNAs can be targeted therapeutically by nucleic acid-based mimics and inhibitors, our findings suggest that modulation of DINO activity may have therapeutic value by restoring TP53 tumor suppressor activity in some tumor types, including HPV positive tumors.

**Figure 9:**
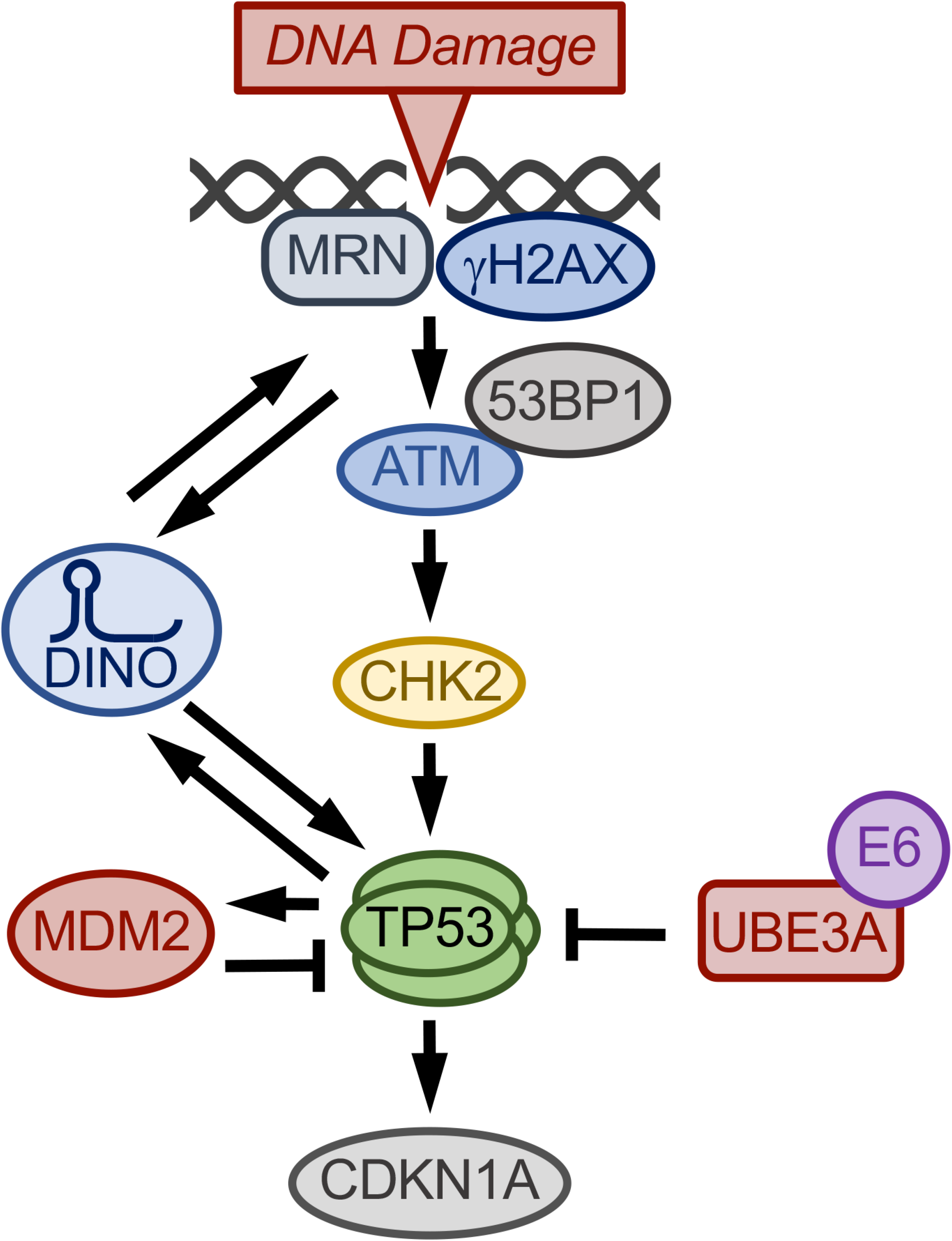
Model of DINO regulation in HPV-positive cervical cancer cells. In response to DNA damage, DINO is induced through an ATM/CHK2/TP53 independent mechanism. DNA damage also triggers TP53 activation through ATM/CHK2 dependent signaling, which causes increased expression of TP53 transcriptional target genes, including CDKN1A and DINO. DINO further activates TP53 by inducing hallmarks of the DNA damage response signaling pathway as evidenced by 53BP1 nuclear foci and ATM and CHK2 activation. By binding to DNA bound TP53 transcription factor complexes, DINO may enhance TP53 transcriptional activity. DINO activity is balanced by TP53-mediated MDM2 upregulation as well as the E6/UBE3A complex, which target TP53 for proteasomal degradation.

## MATERIALS AND METHODS

### Cell culture

Isolation, generation and maintenance of primary human foreskin keratinocytes (HFKs) and HPV16 E6/E7 expressing HFKs has been previously described (44). The C33A, CaSki, HeLa and SiHa cervical cancer cell lines, were purchased from the American Type Culture Collection (ATCC, Manasses, VA) and maintained in Dulbecco’s Modified Eagle Medium (DMEM) supplemented with 10% fetal bovine serum. Subcellular fractionation was carried out using the “Rapid, Efficient And Practical” (REAP) method (77). Doxycycline, cisplatin and doxorubicin were purchased from Sigma-Aldrich (St. Louis, MO). Mitomycin C was purchased from Thermo Fisher Scientific (Waltham, MA). Fluorouracil (5-FU), ATMi (KU-55933) and CHK2i (BML-277) were purchased from Selleck Chemicals (Houston, TX). Acute DINO expression was induced by treating cells with 1 μg/ml doxycycline, made freshly for each use.

### Expression plasmids

The expression vector for acute DINO expression was created by inserting the full-length DINO sequence into the pLIX_403 puro vector backbone (a gift from David Root; Addgene: plasmid #41395). In short, DINO sequences were PCR amplified from an previously published plasmid (a gift from Howard Chang) (43) using primers 5’-CACCGCCGAGTTCCAGC-3’ (forward) and 5’-TTCCAGAATTGTCCTTTATTTATGTCATCTCC-3’ (reverse), and cloned into the Gateway entry vector using the pENTR Directional TOPO Cloning Kit (Invitrogen). DINO sequences were then transferred to the pLIX_403 puro destination vector by standard gateway based in vitro recombination according to the manufacturer’s protocols. The inserted DINO sequences were verified by DNA sequencing. The pLIX-based plasmid for doxycycline regulated green fluorescent protein (GFP) expression was a kind gift from James DeCaprio (78).

### Western blotting and antibodies

Western blots were performed as previously described (44), using TP53 (OP43, Calbiochem), RB1 (OP66, Millipore), CDKN1A (ab109520, Abcam), phospho-ATM (Ser1981) (5833, CST), phospho-Chk2 (Thr68) (2197, CST) and MDM2 (OP46, Millipore), and α-tubulin (ab18251, Abcam) specific antibodies at the dilutions recommended by the manufacturers. Secondary antibodies were horseradish peroxidase (HRP)-conjugated anti-mouse antibody (NA931, GE Healthcare Life Sciences) and HRP-conjugated anti-rabbit antibody (NA934V, GE Healthcare Life Sciences, 1:10,000). Antigen/antibody complexes were visualized by Enhanced Chemiluminescence (Life Technologies) and signals were digitally acquired on a G:Box Chemi-XX6 imager with GeneSys software (Syngene). Protein bands were quantified using GeneTools Software (Syngene).

### RNA interference

The following siRNAs were purchased from Thermo Scientific Dharmacon: non-targeting Pool (D-001810–10), TP53 targeting pool (L-003329-00), and a custom-designed HPV16 E6 siRNA duplex 5’-CCACAGUUAUGCACAGAGCTT-3’ (79). Cells were transfected at a 30 nM siRNA concentration using the Lipofectamine RNAiMax reagent (Invitrogen) per the manufacturer’s instructions.

### Quantitative reverse transcription polymerase chain reaction (qRT-PCR)

Total RNA was isolated and analyzed as previously described (44). The following primer pairs were used for qRT-PCR assays: HPV16 E6: 5’-TGCAATGTTTCAGGACCCA-3’ (forward) and 5’-CATGTATAGTTGTTTGCAGCTCTGT-3’ (reverse); DINO: 5’-GGAGGCAAAAGTCCTGTGTT-3’ (forward) and 5’-GGGCTCAGAGAAGTCTGGTG-3’ (reverse); CDKN1A: 5’-CATGTGGACCTGTCACTGTCTTGTA-3’ (forward) and 5’-GAAGATCAGCCGGCGTTTG-3’ (reverse); RPLP0: 5’-ATCAACGGGTACAAACGAGTC-3’ (forward) and 5’-CAGATGGATCAGCCAAGAAGG-3’ (reverse). The expression data shown was quantified using the ΔΔCT method by normalizing expression of all the qPCR targets to a housekeeping gene, RPLP0.

### Immunofluorescence microscopy

Cells grown on coverslips were fixed in 3.7 % formaldehyde for 30 minutes, permeabilized in 1% Triton X-100 for 10 minutes and blocked in 2.5% serum donkey serum in PBS for 1 hour. After briefly washing in PBS, coverslips were incubated with rabbit anti-53BP1 antibody (ab36823, Abcam, 1:250) for 1 hour at room temperature. An Alexa Fluor-conjugated secondary antibody (donkey anti-rabbit 594, A21207 (Thermo), 1:1000) was used. Nuclei were stained with 4’,6-diamidino-2-phenylindole (DAPI) in ProLong Diamond Antifade Mountant (Thermo Fisher # P36971). Images were captured using a Nikon Eclipse Ti2 microscope. DAPI and Alexa Fluor 594 fluorophores were excited with 405 nm and 594 nm laser lines, respectively. Images with merged DAPI and Alexa Fluor 594 channels were generated using FIJI (https://imagej.net/Fiji).

### Cell viability assays

20,000 CaSki cells per well were seeded in a 12-well plate. Drugs were added at 24 hours post doxycycline induction. At 72 hours after drug treatment, cells were incubated with fresh media containing 10 μg/ml resazurin (Sigma). Resazurin conversion was measured with a Synergy H1 microplate reader (BioTek) with 560 nm excitation and 590 nm emission filters as previously described (80).

## ACKNOWLEDGMENTS

We thank Drs. Howard Chang (Stanford University) and James DeCaprio (Dana Farber Cancer Institute) for generously sharing reagents, Dr. Al Klingelhutz (University of Iowa) for providing telomerase immortalized human foreskin keratinocytes, Dr. Malavika Raman for the use of her fluorescence microscope facility, Warda Aman for help with immunofluorescence experiments, Drs. Amy Yee, Philip Hinds, Peter Juo, Claire Moore and members of the Munger Lab for stimulating discussions and valuable suggestions throughout the course of this work, and the reviewers for their constructive comments and suggestions on this manuscript. Supported by PHS grants AI147176, CA066980 and CA228543 (K.M.) and a Dean’s Fellowship and a Tufts Collaborative Cancer Biology Award from the Tufts Graduate School of Biomedical Sciences (S.S.).

